# Windows out of Africa: A 300,000-year chronology of climatically plausible human contact with Eurasia

**DOI:** 10.1101/2020.01.12.901694

**Authors:** Robert M. Beyer, Mario Krapp, Anders Eriksson, Andrea Manica

## Abstract

Whilst an African origin for Anatomically Modern Humans is well established ^1^, the timings of their expansions into Eurasia are the subject to heated debate, due to the scarcity of fossils and the lack of suitably old ancient DNA ^2^. Here, we estimate potential timings and routes out of Africa by deriving anthropologically and ecologically plausible precipitation requirements for human existence, and applying them to high-resolution palaeoclimate reconstructions for the past 300k years. We find that exit routes and timings previously suggested based on archaeological and genetic evidence coincide precisely with the presence of sufficiently wet corridors into Eurasia, while the gaps between the proposed exit timings co-occur with periods of insufficient rainfall. This demonstrates the key role that palaeoclimatic conditions played for out-of-Africa expansions. The challenging environmental conditions outside of Africa that occurred between windows of potential contact, coupled with the lack of a demographic rescue effect from migration and possible competition with other hominins, likely explain the demise of early colonists prior to the large-scale colonisation of the world beginning from ∼65kya.

## Main text

Analysis of fossil and genetic evidence provides strong support for an African origin of Anatomically Modern Humans (AMHs) ^1^. However, the timing of the out of Africa event has been the focus of recent debate ^2^. The dating of most out-of-Africa fossils, and the dating of the split between Eurasian and African populations based on mitochondrial and whole-genome data ^3,4^ point to a major exit ∼65k years ago. However, archaeological findings in Saudi Arabia dated to at least 85k years ago ^5^, in Israel to at least ∼90k years ago ^6^, possibly ∼194k years ago ^7^, and in Greece dated back to ∼210k years ago ^8^ point to previous excursions from Africa, which might have reached as far as China ^9^. These, or another previous waves, might have also left a small genetic contribution (∼1%) found in modern inhabitants of Papua New Guinea ^10^. Further evidence for a possibly earlier exit comes from traces of geneflow from AMHs into Neanderthal, genetically dated to before 130k years ago, but could be as old as 270k years ago ^11^. Thus, AMHs likely had repeated periods when they were able to leave Africa, even though the timing and fate of those early waves is unclear.

Palaeoclimate reconstructions can provide insights into possible windows out of Africa. Most studies have discussed qualitatively possible scenarios based on a few time slices ^12,13^. Quantitative attempts to define possible windows out of Africa have fitted demographic rules to match either the archaeological record ^14^ or genetic data ^15^. Whilst it is possible to find such rules, it is unclear how biologically realistic they are (e.g. in ref. ^14^, the thermal niche of AMHs changes by 50°C over a period of 125k years). Moreover, the archaeological record is very sparse, especially for the early periods, and genetic data only reflects exits whose descendants have been sampled. Here we take a different approach to characterise suitable periods when AMHs might have left Africa. We use anthropological and ecological data to identify the climatic conditions necessary for AMHs to persist, and then contrast predicted periods of connectivity between Africa and Eurasia with the available archaeological and genetic evidence. We focus on precipitation, as this was the ecologically limiting factor on the regional flora and fauna, and therefore likely the key variable determining the ability of anatomically modern humans to leave the African continent ^16,17^.

To assess connectivity between Africa and Eurasia, we need to establish appropriate precipitation thresholds for areas that would have been suitable to AMHs. We started by inspecting the distribution of contemporary hunter-gatherers from a large anthropological dataset ^18^ (Fig. 1A). Whilst we expect humans to have potentially expanded their geographic range in terms of temperature tolerance, it is less likely that they would have been able to greatly change their ecological niche in terms of aridity. Excluding three populations that are known to reside close to freshwater sources, there is a clear minimum precipitation threshold level at 90 mm of rainfall per year, below which no hunter-gatherers are recorded. This threshold corresponds to a major change in habitat, as it is also the same as the minimum amount of precipitation that can sustain a grazer community ^19^ (Fig. 1A), marking a switch from desert to shrubland ^20^.

**Fig. 1.**
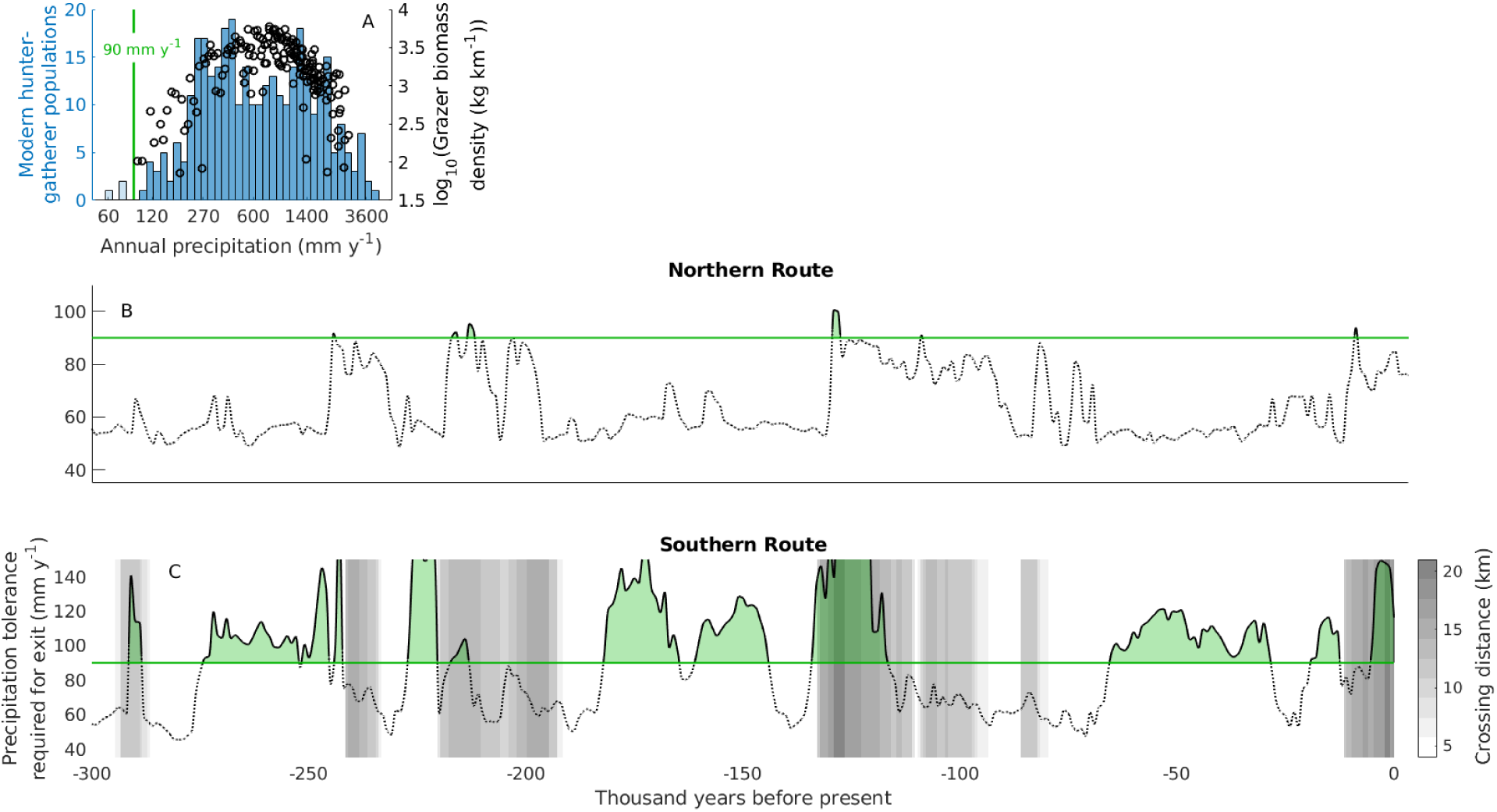
Windows of opportunity out of Africa. (A) Distribution of modern hunter-gatherer populations (blue bins) ^18^ and grazer biomass density (black markers) ^19^ under different precipitation levels. Transparent bins correspond to populations located in immediate vicinity of a water source, which are not considered to be constrained by precipitation. Black lines in (B) and (C) show the minimum precipitation levels that AMHs would have had to withstand at different times to exit successfully along the northern and southern route, respectively. Green lines indicate the critical precipitation tolerance of 90 mm y^-1^. Grey shades in (C) represent the minimum distance needed to cover on water to reach the Arabian peninsula from Africa (Methods).

With this habitability threshold based on anthropology and ecology, we can investigate how compatible inferred periods of migration out of Africa are with the archaeological and genetic record (which were not used in determining the threshold). Until recently, high-quality quasi-continuous palaeoclimate reconstructions were only available for the last 125k years, either from Global Circulation Models such as HadCM3 ^21^, or from simpler Earth Models of Intermediate Complexity such as LOVECLIM ^14^. Here we use a recently developed emulator for the HadCM3 model to generate a high resolution reconstruction of the last 300k years, downscaled to ∼0.5° and bias-corrected to match observed climatic conditions (Methods). We consider the two possible routes out of Africa, through the Nile-Sinai-Land Bridge and the Strait of Bab-el-Mandeb, termed the northern and southern route, respectively ^22^. We started by estimating, at 1k year intervals, the threshold tolerance to low precipitation required to exit through either route; in other words, what was the minimum requirement of annual precipitation that AMHs would have had to withstand to be able to travel all the way out of Africa (Methods). For this estimation, we assumed the Bab-el-Mandeb strait to be always crossable; in reality, the difficulty of this sea voyage would have depended greatly on sea level, which can change the width of the strait from 4 to over 20 km (Methods), and we consider this issue later on in our analysis. Furthermore, we assumed the Nile delta to be crossable at all times irrespective of the local precipitation levels. Modelling other features such as lakes and smaller rivers ^16,23^ over time is challenging; however, given the relatively low threshold suitable of human habitation, we assume that such features would have most likely occurred in areas that received more than the threshold level of 90 mm of rainfall per year.

Based on our ecological threshold, there were only a few windows when the northern route was crossable (Fig. 1B): between 250k and 200k years ago there were a number of peaks in precipitation that came very close or crossed the threshold, and a more consistent period around 130kya, plus a very brief opening around 110k years ago. These latter windows are consistent with archaeological findings in Israel and Saudi Arabia ^5^. After that, this route then likely remained closed until the wet Holocene.

The southern route, on the other hand, was characterised by a number of windows of wetter climate in the southern part of the Arabian peninsula. Based on the precipitation threshold of 90 mm per year, exits through the southern route could have been feasible for a large proportion of the last 300k years (Fig. 1C). Some of these windows of climatic amelioration coincided with very high sea level, when the crossing of the strait would have required a considerable sea voyage ^24^. Prior to the last interglacial, there were three extended intervals of suitably wet climate and low sea level precipitation, from 275k to 242k years ago, from 230k to 221kya, and from 182k to 145kya. During most of the following climatically favourable window (from 135k to 115kya), high sea levels likely impeded migration out of Africa, except at its beginning around 135k years ago. This date coincides with the proposed timing of an early northern exit; thus, if migration did occur, southern migrants might have encountered their northern counterparts on the Arabian peninsula. Following another long period when the southern route was blocked, there was a sizeable window between 65k and 30k years ago, the time that has been long suggested as the main moment of expansion out of the African continent based on both archaeological and genetic evidence. We also note the further connections just after the Last Glacial Maximum, and during the mid-Holocene Eurasian, consistent with evidence of Eurasian backflow into Africa ^25^ (Fig. 1C).

Our reconstructions reveal that there were likely several windows of suitable climate that could have allowed the expansion of AMHs out of Africa, and that the one beginning at 65k, which led to the colonisation of the world by our ancestors, was by no means unique (Fig. 1B,C). Several of these windows predate the earliest remains outside of Africa, but they are entirely compatible with genetic dating of introgression from AMHs into Neanderthal, sometime between 240 and 140k years ago ^11^, and recent dating of material from Israel to ∼170k years ago ^7^ and from Greece to ∼210k years ago ^8^. This suggests that during these earlier windows, AMHs did expand their range towards Eurasia at least once.

A likely explanation of their failure to settle outside Africa is competition with Neanderthal (and possibly Denisovans, whose geographic range is unknown but likely covered a large portion of East Asia ^26^). Competition has also been suggested to have limited the expansion during the last interglacial period ^4,27^, when archaeological evidence points to a more sizeable exit. Our reconstructions (Fig. 1C) suggest that there was an important long period between this window and the following one at 65k years ago when exit from Africa would have been unlikely, thus effectively isolating any of the earlier colonists that might have made it out of their ancestral continent. On the Arabian peninsula, these populations would have faced repeated periods of challenging conditions (i.e. low precipitation) between 115k and 65k years ago (Fig. 2D) that would have led to the likely demise of human populations in that area. In addition, a further expansion would have been predicated on the ability to cross the Taurus-Zagros Mountain range whilst competing with Neanderthal in the north (Fig. 2F) and possibly other hominins, such as Denisova, in the east. With a lack of a demographic rescue effect from further migration out of Africa, most, if not all, remnant populations in Eurasia likely became victims of stochastic local extinctions driven by climatic fluctuations, which significantly affect dispersal success ^28^, and competition during this period prior to the opening of the Africa-Eurasian corridor 65k years ago. The long duration of 35k years of this latter opening (Fig. 1C), combined with technological, economic, social and cognitive changes in AMH societies ^4,27,29^, and possibly the decline of Neanderthal ^30,31^, very probably accounted for the success of the late exit in the subsequent colonisation of Eurasia by AMHs.

**Fig. 2.**
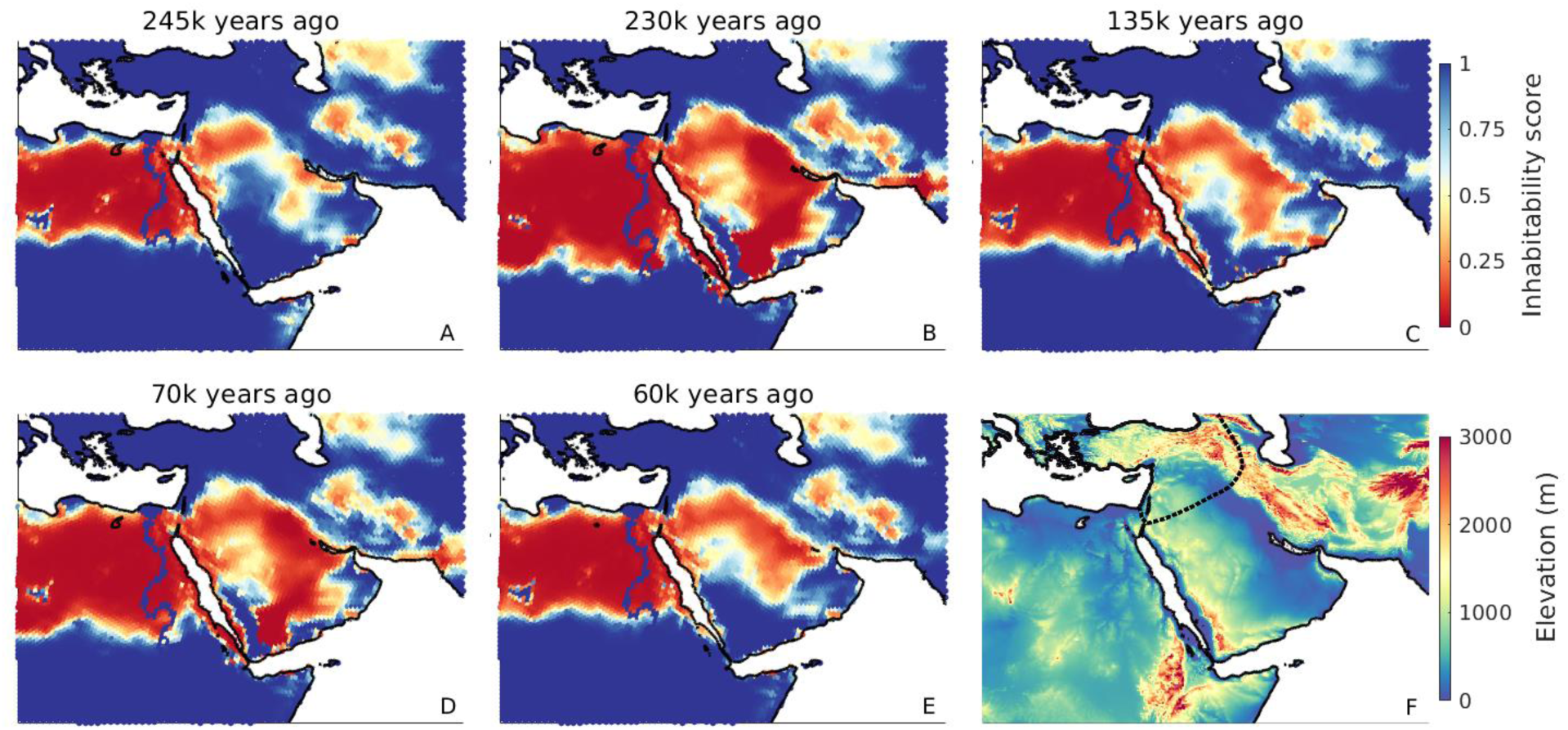
Environmental conditions on the Arabian peninsula. Maps (A)–(E) illustrate precipitation conditions at different points in time. Inhabitability scores are calculated as *I* = (1 + *e*^−*λ*·(*P*−90*mm*)^)^−1^, where *P* denotes local precipitation, and *λ*=0.04 is a smoothing parameter; thus, blue cells with *I*>0.5 represent areas suited for expansions, or climatic refugia during harsh periods, whereas red cells with *I*<0.5 were likely unsuitable for human persistence. (A), (C) and (E) correspond to the estimated timings of northern or southern exits, while (B) and (D) exemplify the challenging conditions between these windows of opportunity. The dotted line in the elevation map (F) represents the known Neanderthal range around 120k years ago ^2^.

## Author contributions

All authors conceived of the study and interpreted the results. M.K. and R.B. generated the palaeoclimate data. R.B. conducted the analysis. A.M. and R.B. wrote the manuscript.

## Supplementary Materials

### Methods

#### Late Quaternary precipitation reconstructions

The accurate reconstruction of precipitation patterns at small spatial scales is crucial for determining the presence or absence of corridors out of Africa, as illustrated in Fig. 2. This requires careful curation of palaeoclimate model outputs, which are typically not available at sufficiently high resolution, and subject to non-negligible biases that can make the difference between a location being represented as inhabitable for AMHs or not. Here we used medium-resolution emulator data for the last 300k years based on precipitation simulations of the HadCM3 climate model. We combined these data with high-resolution HadAM3H simulations generated for selected key time periods, and high-resolution present-era observed precipitation. The downscaling and bias-correction methods used here combine the strengths of each of the three types of input data: the derived precipitation dataset allows for the temporal variability of small scale features, while ensuring values that can be meaningfully compared to ecological data on human precipitation requirements. Climate data obtained by remapping raw climate model outputs, as used in previous studies of human migration ^14,15^, do not share these properties.

Our reconstructions of Late Quaternary precipitation are based on outputs from the global climate emulator GCMET ^32^. GCMET builds on 72 snapshot simulations of the HadCM3 general circulation model ^21,33^, covering the last 120k years in 2k year time steps from 120k to 22k years ago, and 1k year time steps from 21k years ago to the present, at a 3.75°×2.5° grid resolution. On each grid cell, GCMET establishes a local linear regression between the local time series of HadCM3 climate data and four time-dependent forcings (global CO2 and three orbital parameters: eccentricity, obliquity and precession). The values of these predictors are known well beyond the last 120k years; thus, applying them to the grid cell-specific regressions allows to predict continuous global climate up to 800k years into the past. GCMET has been shown to correspond closely to HadCM3 over the last 120k years, and reproduces empirical patterns well in long-term climate reconstructions ^32^.

We used GCMET precipitation data, denoted *P*_*GCMET*_(*t*), covering the last 300k years at 1k year time steps, *t* ∈ *T*_300*k*_. The downscaling and bias-correction approach used here is largely similar to one we previously described in ref. ^34^, and we follow this description here. We downscaled and bias-corrected GCMET precipitation data from its native 3.75°×2.5° grid resolution in two steps (Fig. S1). Both steps use variations of the Delta Method ^35^, under which a high-resolution, bias-corrected reconstruction of precipitation at some time *t* is obtained by applying the difference between lower-resolution present-day simulated and high-resolution present-day observed climate – the correction term – to the simulated climate at time *t*. The Delta Method has been used to downscale and bias-correct palaeoclimate simulations before (e.g. for the WorldClim database), and, despite its simplicity, has been shown to outperform alternative methods commonly used for downscaling and bias-correction ^36^.

A key limitation of the Delta Method is that it assumes the present correction term to be representative of past correction terms. This assumption is substantially relaxed in the Dynamic Delta Method used in the first step of our approach to downscale *P*_*GCMET*_(*t*) to a ∼1° resolution. This method involves the use of a set of high-resolution climate simulations that were run for a smaller but climatically diverse subset of *T*_300*k*_. Simulations at this resolution are highly computationally expensive, and therefore running substantially larger sets of simulations is not feasible; however, these selected data can be very effectively used to generate a suitable (time-dependent) correction term for each *t* ∈ *T*_300*k*_. In this way, we are able to increase the resolution of the original climate simulations by a factor of ∼9, whilst simultaneously allowing for the temporal variability of the correction term. In the following, we detail the approach.

We used high-resolution precipitation simulations from the HadAM3H model ^33^, generated for the last 21,000 years in 9 snapshots (2k year time steps from 12k to 6k years ago, and 3k year time steps otherwise) at a 1.25°×0.83° grid resolution, denoted *P*_*HadAM*3*H*_(*t*), where *t* ∈ *T*_21*k*_ represents the 9 time slices for which simulations are available. These selected data were used to downscale *P*_*HadCM*3_(*t*) to a 1.25°×0.83° resolution by means of the Dynamic Delta Method, yielding

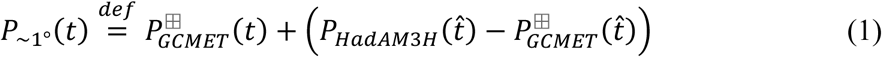

We discuss the choice of an additive approach for all climatic variables in detail later on. The ⊞-notation indicates that the coarser resolution data was interpolated to the grid of the higher-resolution data, for which we used an Akima cubic Hermite interpolant ^37^, which (unlike the bilinear interpolation) is continuously differentiable but (unlike the bicubic interpolation) avoids overshoots. The time 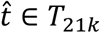 is chosen as the time at which climate was, in a sense specified below, close to that at time *t* ∈ *T*_300*k*_. In contrast to the classical Delta Method (for which 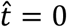 for all *t*), this approach does not assume that the resolution correction term, 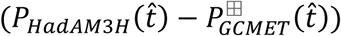, is constant over time. Instead, the fine scale heterogeneities that are applied to the medium-resolution GCMET data are chosen from the wide range of patterns simulated for the last 21k years. The strength of the approach lies in the fact that the last 21k years account for a substantial portion of the glacial-interglacial range of climatic conditions present during the whole Late Quaternary. Following ref. ^32^, we used global CO2, a key indicator of the global climatic state, as the metric according to which 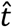 is chosen; i.e., among the times for which HadAM3H simulations are available, 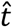 is the time at which global CO2 was closest to the respective value at the time of interest, *t*.

In the second step of our approach, we used present-day (1960-1990) observed precipitation ^38^, *P*_*obs*_(0), to bias-correct and further downscale *P*_∼1°_(*t*) to a hexagonal grid ^39^ with an internode spacing of ∼55 km (∼0.5°). Analogously as before, we obtain

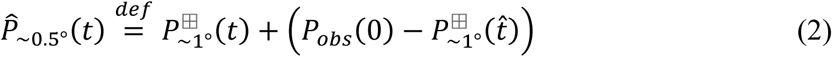

The additive approach used here does not ensure that the derived precipitation is nonnegative across all points in time and space. Thus, we cap values at the appropriate boundaries, yielding

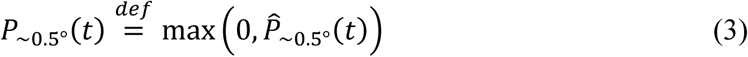

In principle, capping values, where necessary, can be circumvented by using a multiplicative Delta Method that is based on multiplying the relative difference between high- and low-resolution data, at a point in time at which both are available, to the low-resolution data. Instead of Eq. (2), for example, we would have: 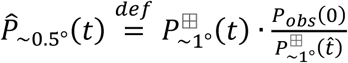. Whilst this approach does ensure non-negative values, it has two important drawbacks that are particularly relevant in arid regions such as the one considered here. First, if present-day observed precipitation in a certain month and grid cell is zero, *P*_*obs*_(0) = 0, then *P*_∼0.5°_(*t*) = 0 in that cell at all points in time, *t*, irrespective of the simulated climate change signal. In particular, this makes it impossible for current extreme desert areas to be wetter in the past. Second, if present-day simulated precipitation in a grid cell is very low (or identical to zero), *P*_∼1°_(0) ≈ 0, then *P*_∼0.5°_(*t*) can increase beyond all bounds. Very arid locations are particularly prone to this effect, which can generate highly improbable precipitation patterns for the past.

We reconstructed land configurations for the last 300,000 years by using present-day elevation ^40^ and a time series of Red Sea sea level ^41^. For locations that are currently below sea level, the Delta method does not work. For these locations, precipitation was extrapolated using a inverse distance weighting approach. With the exception of a brief window from 124-126kya, sea level in the past was lower than it is today; thus, present-day coastal patterns are spatially extended as coastlines move, but not removed.

#### Comparison with empirical proxies

Reconstructions of past wetness and aridity use proxies that reflect not only rainfall conditions but the interaction of precipitation with a series of other local and non-local hydro-climatic variables, such as evaporation and river discharge. As a result, realistic precipitation simulations would be expected to match major qualitative trends of empirical records, rather than exhibit a perfect correlation with the data. Long-term empirical time series of wetness are spatially sparse, particularly in the geographical region considered here. We compared our simulations against three proxy records (Fig. S2). Proxy 1 ^42^ provides a time series of Dead Sea lake levels, for which wet/dry periods are associated with highstand/lowstand conditions (Fig. S2B). Proxy 2 ^43^ represents hydrological conditions at the crossing from Africa to the Arabian peninsula along the southern route, and was obtained from a marine sediment core that allows reconstructing past changes in aridity over land from the stable hydrogen isotopic composition of leaf waxes (dDwax) (Fig. S2C). Proxy 3 ^44^ is an XRF-derived humidity index from a core near the Northwest African coast (Fig. S2D). We find that, overall, the simulated data capture key phases observed in the empirical records for all three proxies well (Fig. S2B–D).

#### Sea level constraints on the southern route

Similar to ref. ^45^, we reconstructed the minimum distance required to cover on water in order to reach the Arabian peninusula (present-day west coast of Yemen) from Africa (present-day east coasts of Eritrea and Djibouti). We used a 0.0083° (∼1km at the equator) map of elevation and bathymetry ^46^ and a time series of Red Sea sea level ^41^ to reconstruct very-high-resolution land masks for the last 300k years. For each point in time, we determined the set of connected land masses, and the distances between the closest points of any two land masses. The result can be graph-theoretically represented by a complete graph whose nodes represent connected land masses and whose edge weights correspond to the minimum distances between land masses. The path involving the minimum continuous distance on water was then determined by solving the minmax path problem whose solution is the path between the two nodes representing Africa and the Arabian peninsula that minimises the maximum weight of any of its edges (Fig. S3).

#### Modelling the expansion of Anatomically Modern Humans

The minimum precipitation value that Anatomically Modern Humans would have to be able to tolerate in order to leave Africa at a given point in time was estimated as follows. For each of the 72 downscaled and bias-corrected precipitation maps representing the last 120,000 years, we initialised populations across northern Africa as shown in Fig. S4. We then tested whether, for an a priori specified precipitation tolerance, there exists a connected path leading out of Africa. This was done by simulating the expansion of populations into neighbouring grid cells if these were above sea level and if the local precipitation level was equal to or larger than the specified precipitation tolerance. If, in the course of this iterative process, a grid cell east of 60.2° longitude or north of 42.5° latitude was populated, then the tested precipitation tolerance was considered sufficient for a complete out of Africa exit (Fig. S2a). If, otherwise, the target longitude or latitude thresholds were not crossed by the time no new cells are populated, then the tested precipitation tolerance was considered insufficient to leave Africa (Fig. S2b). Starting from precipitation tolerances of 1000 mm y^-1^ and 0 mm y^-1^, for which leaving Africa is possible at all times and at no time, respectively, we iteratively converged to the critical precipitation tolerance with a precision of 0.1 mm y^-1^ by means of the bisection method. In order to specifically determine the precipitation tolerance needed for the northern and southern exit, we rendered the passage of the respective other route impassable (Fig. S2).

## Supplementary Figures

**Fig. S1.**
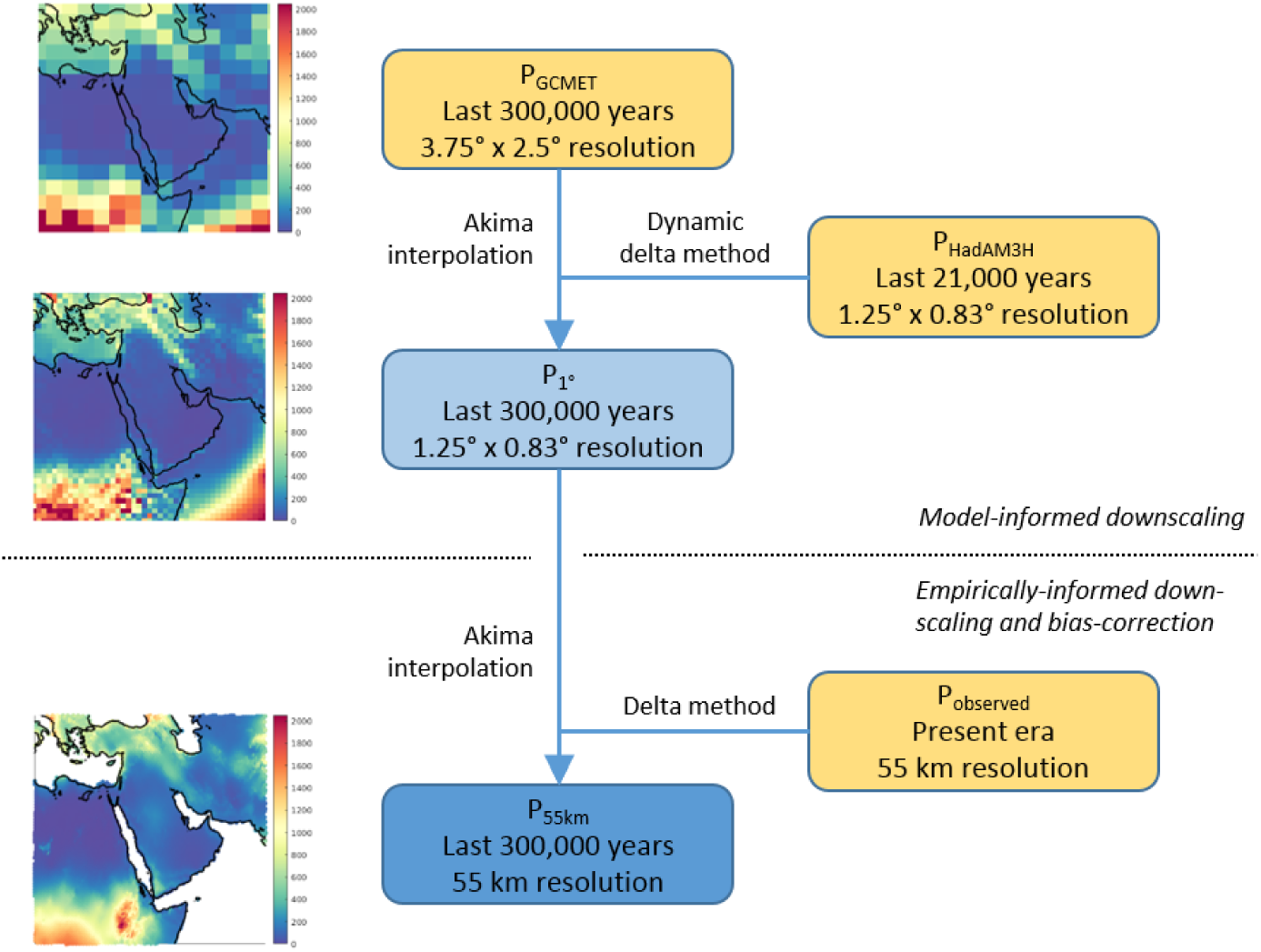
Reconstructing high-resolution bias-corrected precipitation for the Late Quaternary. Yellow boxes represent raw simulated and observed data, the dark blue box represents the final dataset. Maps correspond to the datasets represented by the left three boxes.

**Fig. S2.**
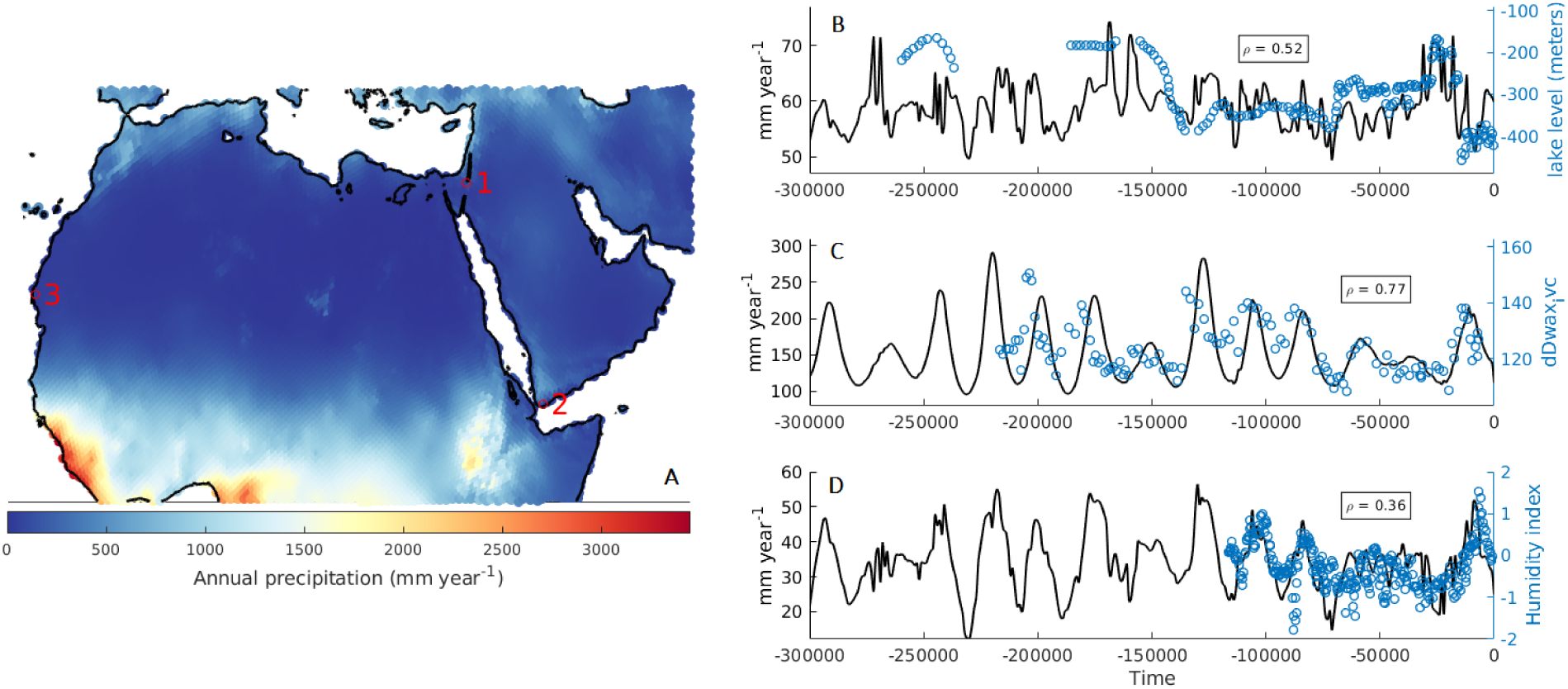
(A) Geographical locations of empirical proxies shown on a present-day precipitation map. (B)–(D) Comparisons of simulated precipitation against the three empirical wetness proxies over time, and the corresponding Pearson correlation coefficients ρ.

**Figure S3.**
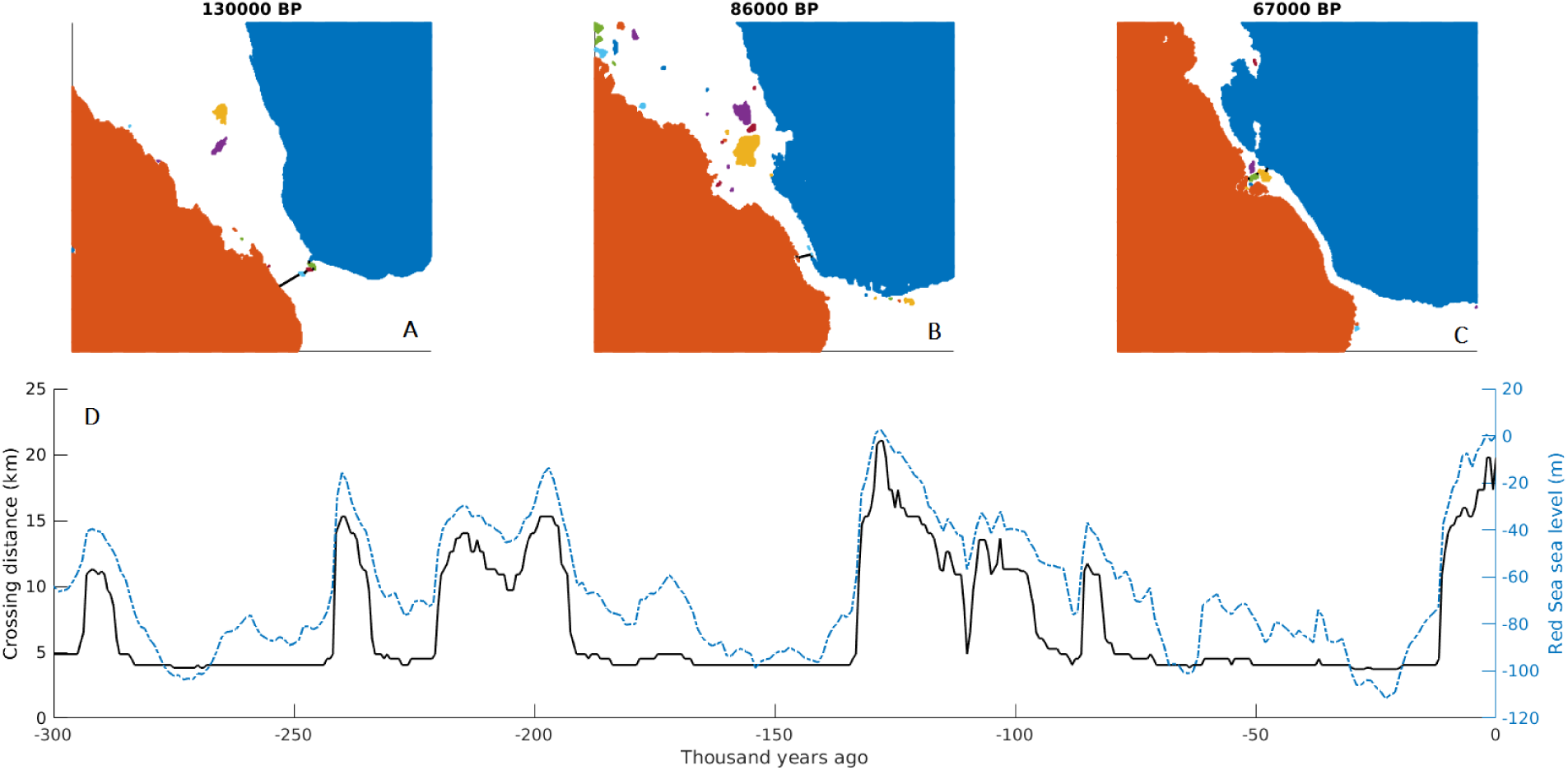
Minimum crossing distance between Africa and the Arabian peninsula. (A)-(C) show reconstructions of the land configuration at selected times based on Red Sea sea level (blue line in (D)), with colours indicating separate land masses. Black lines in (A)-(C) represent the path connecting Africa and the Arabian peninsula that minimises the distance required to continuously cover on water. The black line in (D) shows this distance for the complete time period.

**Fig. S4.**
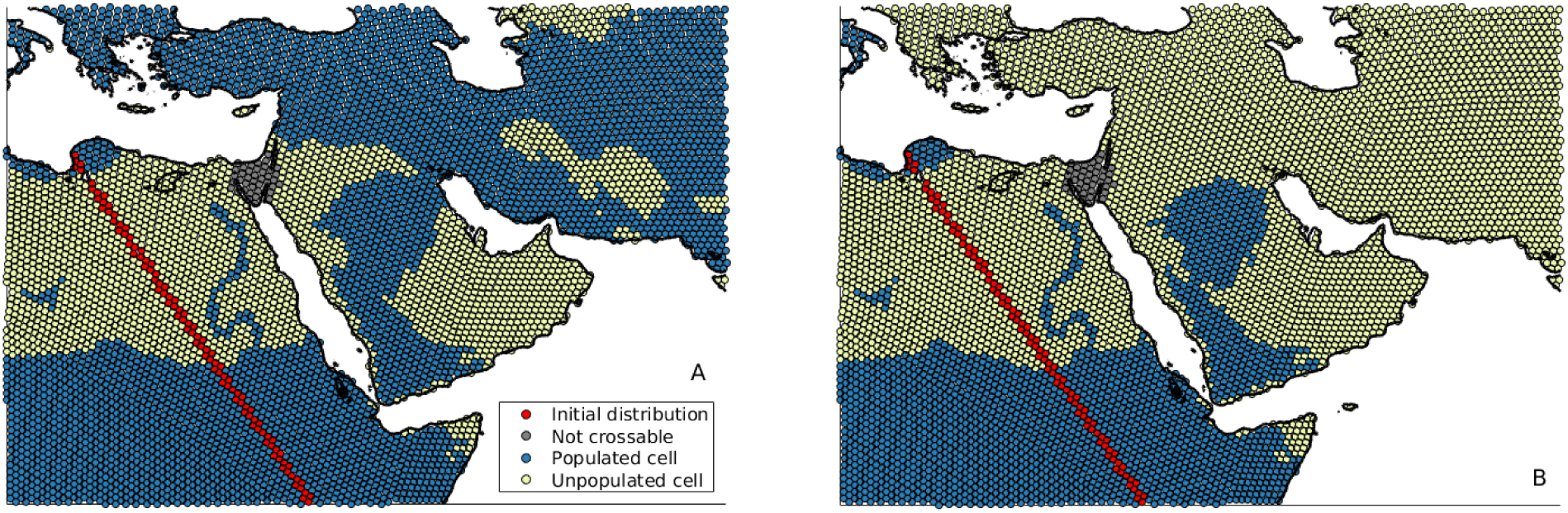
Terminal (A) successful and (B) failed southern exits simulated for precipitation tolerances of 110 mm y^-1^ and 120 mm y^-1^, respectively, and an underlying present-day precipitation map. In the simulation shown, the northern Route was rendered uncrossable.

